# Emergence of carbapenem, beta-lactamase inhibitor and cefoxitin resistant lineages from a background of ESBL-producing *Klebsiella pneumoniae* and *K. quasipneumoniae* highlights different evolutionary mechanisms

**DOI:** 10.1101/283291

**Authors:** Eva Heinz, Hasan Ejaz, Josefin Bartholdson Scott, Nancy Wang, Shruti Guanjaran, Derek Pickard, Jonathan Wilksch, Hanwei Cao, Ikram ul-Haq, Gordon Dougan, Richard A Strugnell

**Author notes:** Correspondence: R.A. Strugnell, Department of Microbiology and Immunology, The University of Melbourne, at Peter Doherty Institute for Infection and Immunity, Melbourne, Australia; Eva Heinz, Infection Genomics Program, Wellcome Trust Sanger Institute, Hinxton, CB10 1SA, UK.

## Abstract

*Klebsiella pneumoniae* is recognised as a major threat to public health, with increasing emergence of multidrug-resistant lineages including strains resistant to all available antibiotics. We present an in-depth analysis of 178 extended-spectrum beta-lactamase (ESBL)-producing *Klebsiella* strains, with a high background diversity and two dominant lineages, as well as several equally resistant lineages with less prevalence. Neither the overall resistance profile nor the virulence factors explain the prevalence of some lineages; we observe several putative hypervirulence factors across the population, including a reduced virulence plasmid, but this does not correlate with expansion of one or few highly virulent and resistant lineages. Phenotypic analysis of the profiles of resistance traits shows that the vast majority of the phenotypic resistance profiles can be explained by detailed genetic analyses. The main discrepancies are observed for beta-lactams combined with beta-lactamase inhibitors, where most, but not all, resistant strains carry a carbapenemase or *ampC*. Complete genomes for six selected strains, including three of the 21 carbapenem-resistant ones, are reported, which give detailed insights into the early evolution of the *bla-NDM-1* enzyme, a carbapenemase that was first reported in 2009 and is now globally distributed. Whole-genome based high-resolution analyses of the dominant lineages suggests a very dynamic picture of gene transfer and selection, with phenotypic changes due to plasmid acquisition and chromosomal changes, and emphasize the need to monitor the bacteria at high resolution to understand the rise of high-risk clones, which cannot be explained by obvious differences in resistance profiles or virulence factors.

**Importance:** Carbapenem-resistant and extended-spectrum beta-lactamase (ESBL) carrying *Enterobacteriaceae* were recently highlighted as critical priority fo the development of new treatments by the WHO. *Klebsiella pneumoniae* is a member of the *Enterobacteriaceae* and has seen a dramatic rise in clinical relevance due to its uncanny ability to accumulate multidrug-resistance plasmids. We present a detailed analysis of a set of ESBL-resistant *K. pneumoniae* clinical isolates, and our high-resolution whole-genome sequence analyses highlight that acquisition of drug resistances is not a one-way street in *K. pneumoniae*, but a highly dynamic process of gain and loss, and that the most successful lineages in the clinic are not necessarily the most resistant or most virulent ones. Analysis of the virulence potential also shows that these strains harbour some, but not all, hallmarks of hypervirulent strains, emphasizing that it is not a clear distinction between hypervirulent and other strains, but equally in flux.

## Introduction

The past four decades have seen a continuous escalation of bacterial pathogens acquiring resistance mechanisms against antimicrobials, and especially antimicrobial resistance determinants associated with mobile elements have spread with exponentially increasing speed across the globe (1, 2). A particularly successful pathogen in this group is *Klebsiella pneumoniae*, which was formerly known as a major cause of infections in neonates, especially in developing countries (3–6) and community-acquired and nosocomial infections in immunocompromised patients (7–9). The acquisition of extended-spectrum beta-lactamases (ESBLs) rapidly increased in *Klebsiella* spp. from the 1990s, particularly in hospital isolates (10, 11). In 2009 there was first description of NDM-1, a metallo-betalactamase which hydrolyzes carbapenems and can escape beta-lactamase inhibitors, which were until then the main alternative treatments for third-generation cephalosporin-resistant bacteria (12). NDM-1 appears to have originated as a chromosomal gene fusion in *Acinetobacter* (13), but rapidly disseminated around the globe as well as across several species (e.g. *E. coli*, *Provatella;* 14).

We report a high-resolution analysis of a set of ESBL-carrying *K. pneumoniae* isolates from a tertiary care children’s hospital in Lahore, Pakistan, collected between 2010 and 2012 (15). The selected strains were all ESBL producers, and our population study shows that whilst we observe two dominant lineages, no clear advantage for these lineages can be seen either in the resistance profile, or the virulence factor repertoire, despite some strains carrying markers of hypervirulent *Klebsiella;* these markers were not widely disseminated within the dataset, but occurred intermittently.

The main treatment alternatives for ESBL-producing *Enterobacteriaceae* are i) cefoxitin, which is resistant to ESBL enzymes but sensitive to AmpC; ii) beta-lactams in combination with beta-lactamase inhibitors, with two widely used combinations piperacillin/tazobactam and amoxicillin/clavulanic acid; and iii) carbapenems. Comparing phenotypic with predicted resistance profiles showed that resistance to carbapenems correlated with the presence of carbapenemase. Resistance against either or both of the alternative treatments (cefoxitin or the use of beta-lactamase inhibitors) could be explained in some cases through the presence of NDM-1, whereas for several strains, the resistance mechanisms are still unclear.

We furthermore have added complete genome sequences of six selected strains including one representative for each lineage harbouring the *bla-NDM-1* gene; this gives us important insights into the early spread of carbapenemases and the associated plasmid diversity. We note no differences in the *bla-CTX* or *bla-TEM* genes, but high variation in allelic distribution of *bla-OXA* and *bla-SHV*, including several ESBL variants and additional plasmid-encoded copies of *bla-SHV*. Our study highlights the mobile and plastic nature of *Klebsiella* resistance determinants, which vary widely, even within strains selected for similar resistance traits from a single hospital, and within the same sequence type. This dynamic relationship between a now key opportunistic pathogen and a very large pool of mobile and readily transferrable resistance mechanisms is very sobering and is in stark contrast to observations of other resistant pathogens, where acquisition of an advantageous resistance/mobile element leads to rapid, clonal spread of a distinct lineage linked to the resistance element (16–18).

## Results

### Strain collection shows high variety of sequence types and virulence determinants

The strains were collected between 2010-2012 through the routine microbiological screening in The Children’s Hospital, Lahore, Pakistan, and were pre-selected for ESBL presence through E-test (15). The population structure of our dataset includes a high diversity of sequence types (15; Fig. 1A, 1B), as well as a high number of isolates from *K. quasipneumoniae*, a subspecies that was previously thought to be less virulent (Fig. 1; 15). In contrast, no *K. variicola* was identified, a closely related species which is also routinely identified as *K. pneumoniae* with non-distinguishable clinical symptoms. To analyse the molecular epidemiology in finer detail, we identified the capsule (K) and lipopolysaccharide O-antigen types based on their respective operons in the genomes, which are important markers to monitor the epidemiology of *Klebsiella* spp. Recent work (19, 20) highlighted the considerable diversity found among different *Klebsiella* capsule types (21), and this is also reflected in the collection reported here, where the diversity of sequence types and capsule types is equivalent, with a different capsule type for almost each sequence type (Fig. 1C, Table S1). In contrast to capsule loci, there is less diversity of LPS O-antigens with two main types in our collection; however these are not the usually reported main types (i.e. not O1, O2 and O3; 20, 22) but include less common types. LPS serotype 05 was the second most common, present in 68 of our isolates across 3 sequence types (Fig. 1).

**Fig. 1:**
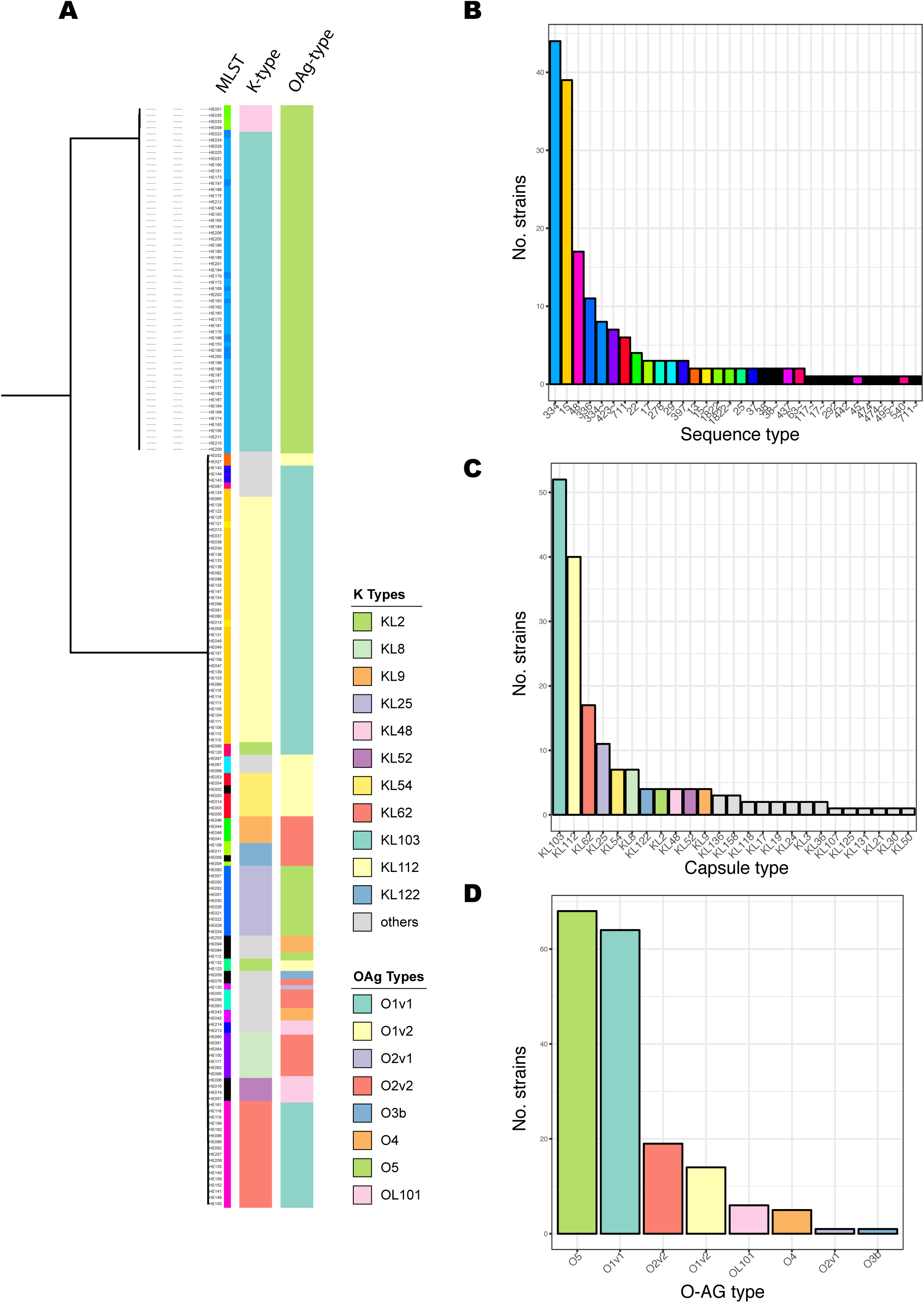
LPS O-antigen, capsule and sequence types in this study. **(A)** The guide tree is based on roary as in Fig. 1; and shows the diversity of capsule and O-antigen type, which correlate with sequence types (one sequence type usually shares the capsule- and O-antigen combination). **(B)** Two main (ST15, ST334) sequence types dominate the collection, followed by several medium-abundance (e.g. ST45, ST336) types and a high diversity of low-abundance sequence types. **(C)** The capsule types almost share almost exactly the same level as diversity as the sequence types, with a different capsule type associated with sequence types; whereas the O-antigen types **(D)** are as expected less diverse, but dominated through the commonly as less diverse perceived type O5.

**Table 1:** Detection of carbapenemase using minimum inhibitory concentration and gene prediction (ariba read coverage).

| Strain | MIC (Meropenem) | NDM-1 coverage |
| --- | --- | --- |
| HE006 (14893_8#74) | 64 | 75.0 |
| HE007 (14893_8#75) | 32 | 83.8 |
| HE021 (14893_8#89) | 2 | Not Detected |
| HE016 (14893_8#84) | 64 | 59.3 |
| HE019 (14893_8#87) | 64 | 79.1 |
| HE121 (14936_2#2) | 32 | 105.4 |
| HE122 (14936_2#3) | 16 | 49.4 |
| HE065 (14936_3#41) | <1.25 | Not Detected |
| HE125 (14936_2#6) | 32 | 73.3 |
| HE128 (14936_2#9) | 32 | 43.4 |
| HE001 (14893_8#69) | 16 | 8.1 |
| HE035 (14936_3#11) | 4 | Not Detected |
| HE063 (14936_3#39) | 16 | 6.2 |
| HE055 (14936_3#31) | 16 | Not Detected |
| HE060 (14936_3#36) | 64 | 44.7 |
| HE061 (14936_3#37) | 64 | 47.1 |
| HE062 (14936_3#38) | 64 | 53.8 |
| HE213 (14936_2#94) | <1.25 | Not Detected |
| HE064 (14936_3#40) | 64 | 52.9 |
| HE066 (14936_3#42) | 64 | 56.3 |
| HE100 (14936_3#76) | 64 | 43.5 |
| HE117 (14936_3#93) | 64 | 53.9 |

With the collection of strains there is an almost ubiquitous presence of heavy metal resistance (silver, *sil*; copper, *pco*) as well as the type 3 fimbriae (i.e. *mrk*; Fig. S1). The genes encoding the siderophores aerolysin and yersiniabactin are distributed only partially throughout the collection (Fig. S1; 23), and no salmochelin or colibactin encoding genes were identified. Our collection encodes for six different types of yersiniabactin in eight sequence types (23), including the less common combination of ybt8 with ICEKp3 (ST423), and one sequence type (ST48) with three different variants (Fig. S1), highlighting the wide distribution of the different siderophore types even within a confined setting already in these historical isolates.

We identified a high number of genes predicted as pseudogenes by ariba (Fig. S2A), including the virulence gene *rmpA2*, the capsule upregulator (Fig. S2B), as well as the fluoroquinolone resistance gene *qnrB* (Fig. S2C). RmpA is inactivated through a frameshift in either the poly-G and poly-A regions (Fig. S2B), as described previously (24). RmpA and aerobactin are indicative of the presence of the *Klebsiella* virulence plasmid (25–28), and we therefore compared several virulence plasmids found with our data, and we performed comparisons against the plasmid pSGH10, which is present in a representative model strain of the hypervirulent clade CG23 (28). The plasmid is not evenly spread across the sequence types from Lahore and lacks salmochelin; both when compared mapping (Fig. 2A) or assemblies (Fig. 2B). The presence or absence of these putative key virulence factors (15) could not be correlated with clinical outcome (details Table 2).

**Fig. 2:**
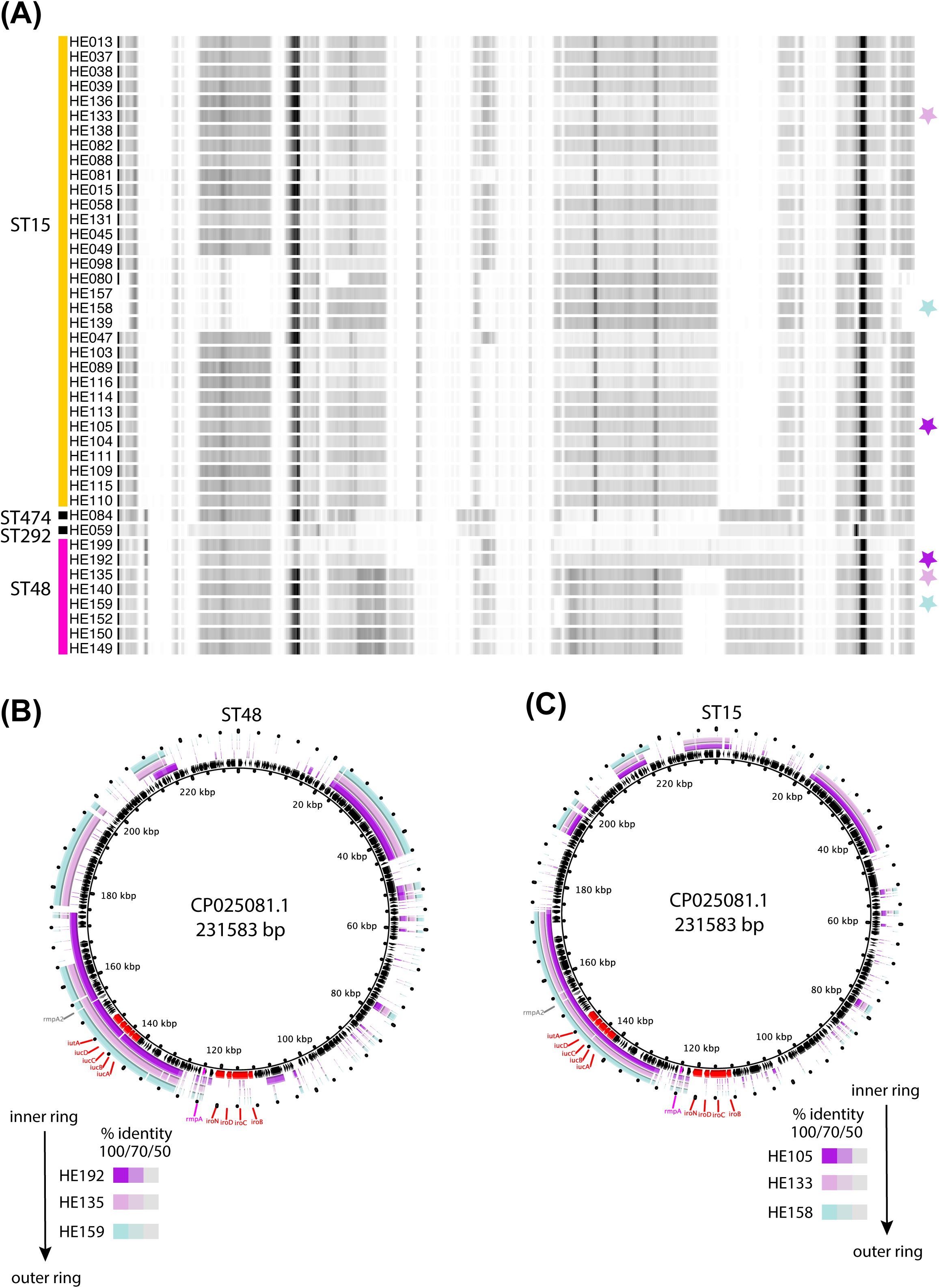
Comparison of strains with *rmpA2* pseudogenes to Klebsiella virulence plasmids. **(A)** shows a heatmap from mapping the reads against the recently published reference plasmid for hypervirulent strains, pSGH10 (28). Representative strains were chosen as indicated by star icons in **(A)** for ST48 **(B)** and ST15 **(C)**, the relevant Illumina contigs extracted using blastn and abacas, and mapped against the reference plasmid using BRIG (97) to further illustrate the partial conversation of the virulence plasmid also in the assembled data.

**Table 2:** Testing outcome versus presence of yersiniabactin or aerobactin.

|  | outcome |  |  |
| --- | --- | --- | --- |
| *Siderophore* | Death | Discharged | Total |
| *Yersiniabactin* + | 16 | 26 | 42 |
| *Yersiniabactin* - | 34 | 75 | 109 |
| *Total* | 50 | 101 | 151 |
| *p-value*<sup>1</sup> | $p_{chi}=0.5388; p_{fisher}=0.4444$ | | |
| *Aerobactin* + | 13 | 22 | 35 |
| *Aerobactin* - | 37 | 79 | 116 |
| *Total* | 50 | 101 | 151 |
| *p-value*<sup>1</sup> | $p_{chi}=0.709; p_{fisher}=0.6823$ | | |

<sup>1</sup>: P-value determined via a chi-square test of independence using the R function `chisq.test(input matrix)` annotated as $p_{chi}$ ; and a fisher's exact test using the R function `fisher.test(input matrix)` annotated as $p_{fisher}$ .

Most multidrug-resistance in *Klebsiella* is caused by acquired resistances on mobile elements, which is clearly reflected in our dataset (Fig. S1), with the expected high number of resistances against possible first-line drug classes aminoglycosides, fluoroquinolones, sulphonamides and beta-lactams, noting that the isolates were selected for ESBL expression, as well as other antimicrobials (Fig. S1).

### Comparison of predicted to measured MICs highlights importance of genetic background and potential unknown resistance determinants

Measured MIC values of widely-used antimicrobials of our strain collection generally matched the genomics-predicted resistances (Table S3, Fig. S3); intrinsic resistance mechanisms such as deactivation of porins, upregulation of efflux pumps or resistance genes, might account for some of the non-matching results (29–31). Resistance against aminoglycosides is conferred almost exclusively by AAC3 or AAC6 (found in 87% of all isolates); however, it is noteworthy that despite some of the widespread key integrons in *Klebsiella* spp. usually including aminoglycoside resistance (32, 33) there is still sensitivity against this class in our group of clinical isolates. Thus, this antibiotic class might still provide a treatment option for some ESBL-positive strains.

Closer investigation of the potential loss of fluoroquinolone resistance via the predicted *qnrB* pseudogenes (Fig. S2A) showed that it might reflect a functional gene with an alternative start codon (Fig. S2C), which however is recognised as truncated due to its slightly shorter sequence at the N-terminus. The full-length sequence found in the CARD and ARGANNOT database (accession ABC86904.1) contains 12 amino acids at the N-terminus. A frameshift mutation present in the *qnrB* coding sequences of our strain set (Fig. S2A) does not affect the sequence after the alternative start codon. Comparison to the structural information of the functional QnrB (34) shows that the N-terminal structural elements are conserved despite the shorter N-terminus (Fig. S2C; coloured boxes indicate structural elements as in Figure A1 from 34); the remainder of the predicted proteins in our data are 100% identical with the reference sequences. We also note that the sequence in Genbank has been updated (ABC86904.2), which renders it the same length as the more recently described qnrB2 (35; APU91821.1, 5 amino acids difference), and we therefore assume the putatively shortened QnrB to be functional and to confer quinolone resistance in the strains reported here. Only few lineages show additional resistance to fluoroquinolones through mutations in *gyrA* and *parC*.

### Alternative beta-lactam treatments for ESBL-positive organisms

*Klebsiella* spp. harbour two intrinsic enzymes related to low-level beta-lactam resistance; AmpH, an AmpC-related enzyme functioning as penicillin-binding protein; and a chromosomally integrated beta-lactamase of the SHV, OKP or LEN family for *K. pneumoniae*, *quasipneumoniae* and *variicola*, respectively (36). In addition to these chromosomal enzymes, we detected CTX-M-15, three OXA and three TEM variants which however result in the identical amino acid sequence, VEB-5; the AmpC enzymes CMY-6 and CMY-18, and NDM-1 carbapenemases as well as additional copies of SHV (Fig. 3).

**Fig. 3:**
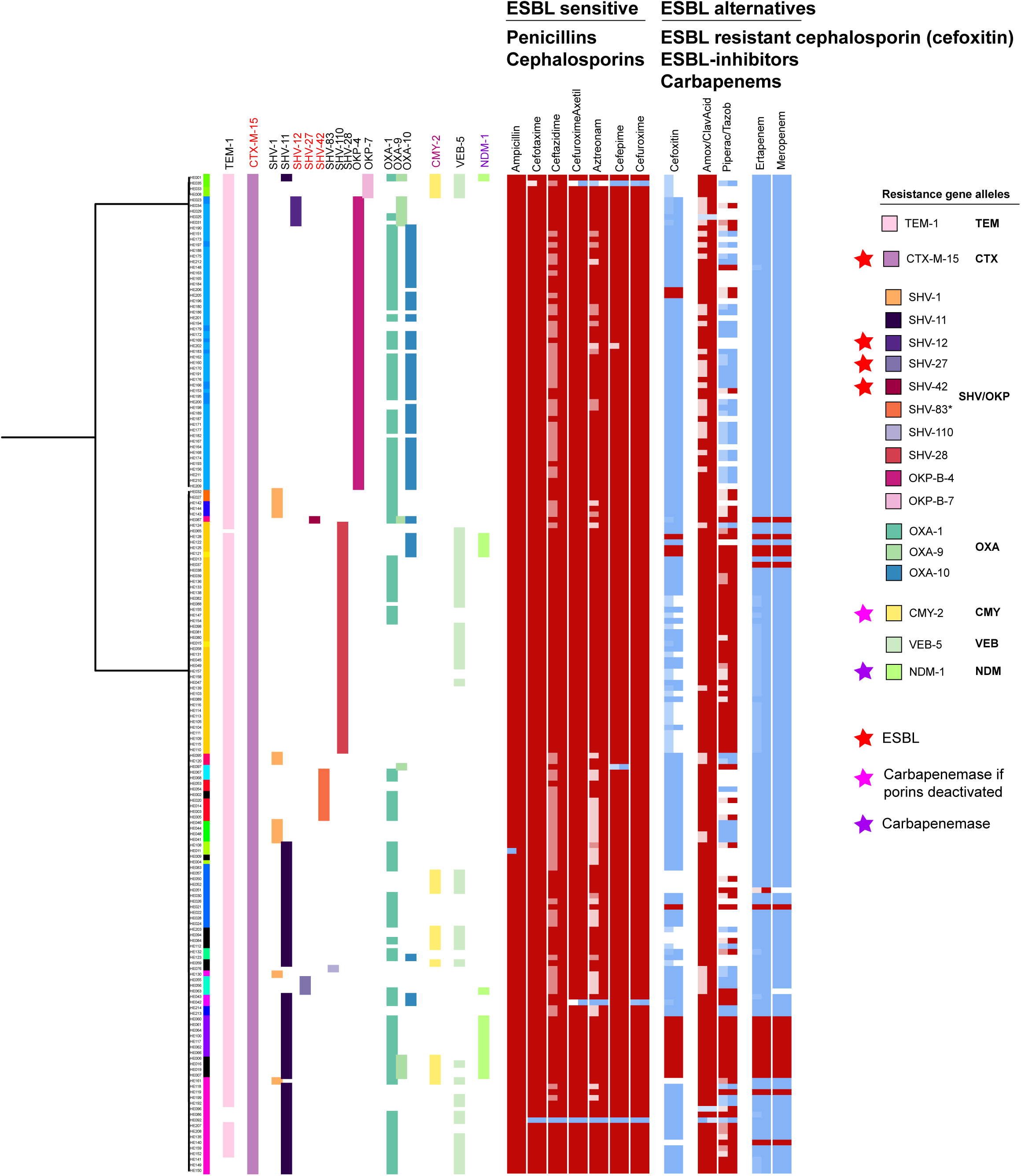
Details of predicted beta-lactamase enzymes and resistance profiles to treatment alternatives for ESBL organisms. (**A**) The guidance tree is as in Fig. 1. All strains with TEM or CTX genes encode for the broad-spectrum beta-lactamase TEM-1 and the ESBL enzyme CTX-M-15, respectively, and only the VEB-5 and NDM-1 allele, and two alleles for the AmpC CMY (6 and 18) could be identified. On the contrary, we see (amino acid) variation in the different alleles for SHV/OKP and OXA; colours are according to **(B)** for SHV/OKP and **(C)** for OXA. *Bla-*SHV can have different activity spectrums; the red stars indicate extended-spectrum beta-lactamase activity. Pink indicates weak carbapenemase activity which needs additional mutations (e.g. porin deactivation) to confer resistance, dark violet denotes carbapenemase activity. The allele assignment was controlled at the beta-lactamase online database (http://www.laced.uni-stuttgart.de); activity assignments are according to (98).

Sensitivity was observed among a high number of isolates for cefoxitin and piperacillin-tazobactam; this is expected as cefoxitin is generally insensitive to ESBL (e.g. *bla-CTX-M-15*, which is present in all but one of our isolates), whereas resistances against piperacillin are widespread, but the beta-lactamase inhibitor tazobactam is still usually effective. Resistance against cefoxitin can be explained in most cases by the presence of the carbapenemase *bla-NDM-1* which hydrolyses cefoxitin, the *bla-CMY* AmpC, or both; only three strains (HE021, HE205, HE206; Fig. 3) seem to confer resistance due to other factors. A pan-genome analysis of the *K. quasipneumoniae* lineage reveals a number of differences which might tangentially affect drug sensitivity such as transposases and hypothetical proteins (likely phage-derived following comparisons to public databases), however the only clear genetic difference between HE205 and HE206 is an iron uptake system (*fec*) that is absent from the two former strains (Fig. S4).

It is recognised that sensitivity against piperacillin/tazobactam (TZA) often coincides with lower MIC values against amoxicillin/clavulanate (Fig. S4; Table S2), even though, based on a phenotypic sensitive/resistant evaluation, these strains are assessed as resistant. More detailed analysis of the encoded SHV/OKP *bla* genes revealed that several *K. quasipneumoniae* strains (HE031, HE029, HE025, HE034, HE023; ST334) gained a *bla-SHV* gene in addition to the intrinsic *bla-OKP*, and the contigs encoding the *bla-SHV* copies are similar to plasmid sequences based on Blast data from public websites, and a plasmid location is also supported by the unique plasmid replicon profile of these strains (Fig. S1). These putatively plasmid-borne *blaSHV* genes encode for a sequence with the mutations G238S and E240K, which correlate with TZA resistance, or TZA-intermediate sensitivity (Fig. 3; 37–39). We also observed several sequences in our dataset encoding the L35Q mutation; however this only provides a subtle increase in TZA MICs (37, 40, 41), and no mutation was found at S130 (42).

### The emergence of NDM-1 on mobile elements

The MIC data highlights that the same gene (NDM-1) might cause different resistance phenotypes in different genetic backgrounds (Table 1). This could be due to both the genetic background provided by the bacterial chromosome, as well as the plasmid harbouring the gene. To identify the number of different plasmids that were present in our dataset, and to test how many of them are still circulating as reported in recent literature, we performed PacBio sequencing of representative isolates (Table 1, S1).

Since some NDM-1 genes detected in Illumina reads could not be confirmed by PacBio sequencing, and were only present in low coverage (Table 1), we performed additional MIC tests for meropenem (E-tests, see methods; Table 1). This confirms that the low-coverage enzymes only resulted in lower MICs read as sensitive, which can indicate either loss of the plasmid carrying NDM-1, or of a highly mobile cassette carrying NDM-1. The unstable nature of NDM-1 has been observed previously in several independent reports, where loss can occur already after two generations of re-growth, and even under meropenem selection (43–45). The loss of the entire plasmid carrying NDM-1 in part of our samples, or at least a larger transposable cassette, is further supported by the presence of several resistance genes predicted at low abundance and absent in the PacBio assemblies, potentially indicating the unstable nature of the plasmid carrying the resistance (Fig. 4).

**Fig. 4:**
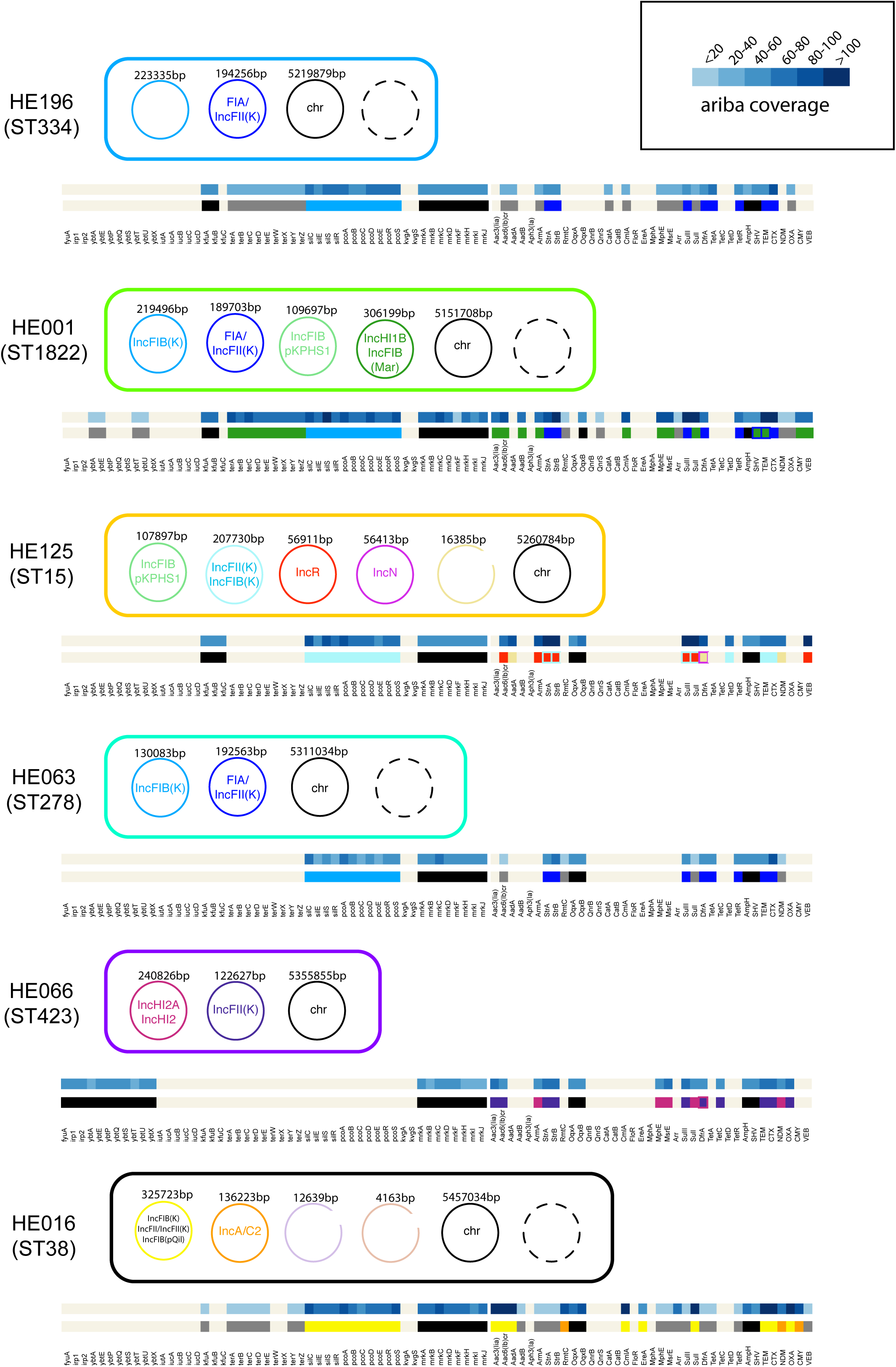
The plasmid diversity for representative isolates. The contigs obtained from PacBio for selected strains are shown; plasmid replicons as predicted by the PlasmidFinder web server (https://cge.cbs.dtu.dk/services/PlasmidFinder/, 94). Below each strain are the resistance genes as predicted by ariba on a colour gradient displaying the read coverage of the hit; and the row below these indicate on which plasmid the respective genes were found (colours according to the respective cell overview). Black indicates chromosomal, dark grey indicates gene only found in Illumina reads but not in the PacBio data. The presence of genes was investigated by blastn, only complete perfect (100% identity) hits were considered.

In addition, replicon analysis and mapping of the resistance and virulence genes back to the plasmids across all strains where the genomes were resolved with PacBio indicates the involvement of several different plasmids, showing highly dynamic profiles between strains even within closely related isolates (Fig. 4). Almost fixed in the population across all sequence types are as expected the incompatibility groups F and H; with only few isolates showing rarer types such as N and R (Fig. S1, Fig. 4).

Two of the plasmids show the original, more complete cassette including the GroEL/S from the original *Acinetobacter* construct (Fig. S5), whilst in one case a smaller portion of this gene region is conserved; this variation in the different forms of NDM-1 has been described in other studies (46, 47).

### Comparison ST15 isolates from Lahore with outbreak in Nepal

To gain further insights into the diversity within dominant lineages in Fig. 1, we analysed the differences between the two ST15 lineages reported in Nepal (48, 49), and in this study (Table S3). This included assessing the presence or absence of genes via a pan-genome analysis of a subset of isolates, as well as mapping of the short read data for the same isolate against a PacBio-sequenced reference sequence (accession number CP008929; Fig. S6, Fig. 5), derived from the Nepal outbreak which has the complete genome (49). A comparison of the plasmid content showed entirely different compositions, with only one plasmid that was commonly conserved (Fig. S7). Analysing the capsule operons, we observed the same novel capsule type as described for the Nepal isolates (48; Fig. S6); further highlighting the likely close relationship of this virulent lineage that seems to be spreading throughout this area of the world.

**Fig. 5:**
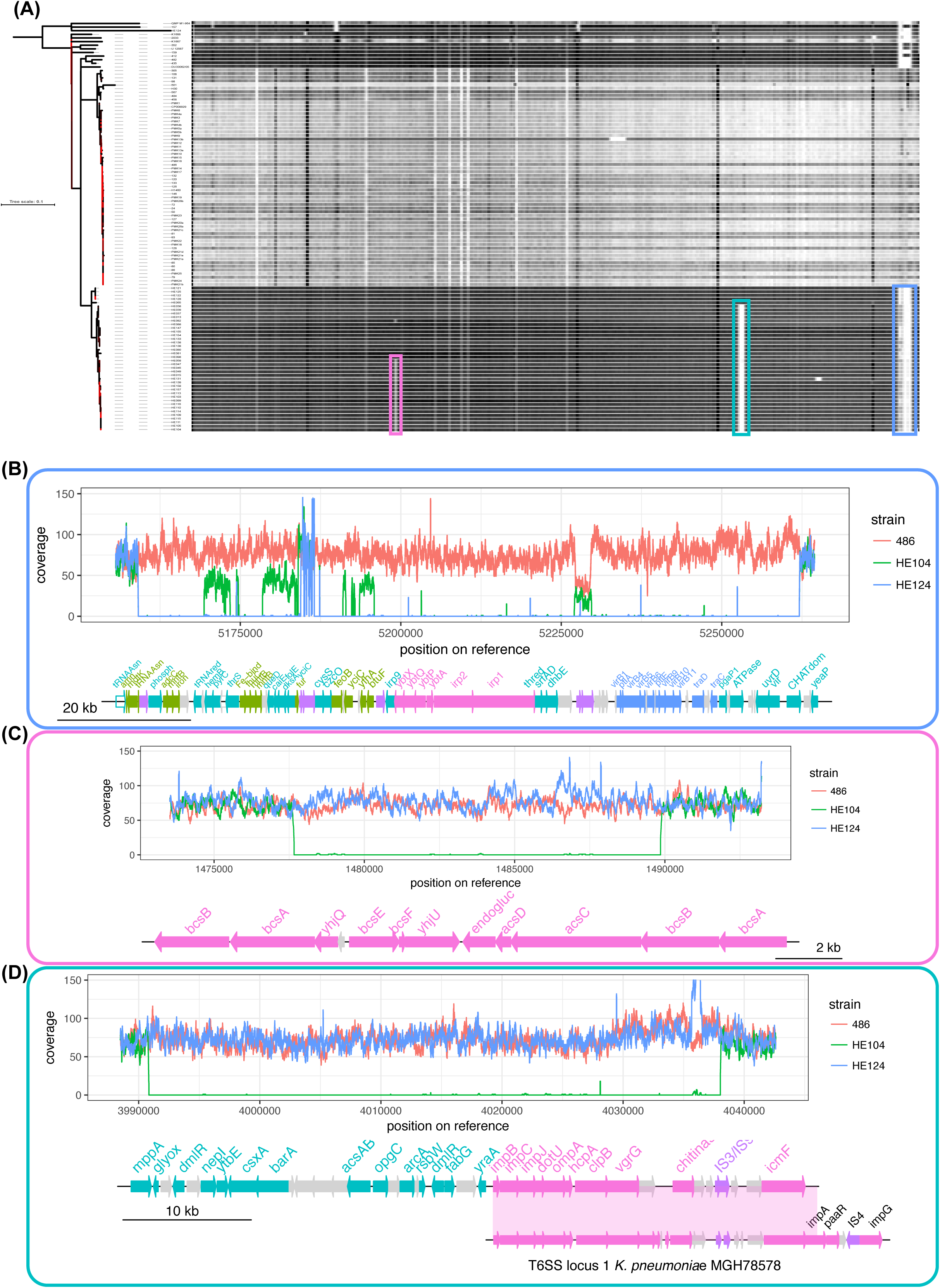
In-depth comparison of expanded lineages. **(A)** The strains belonging to ST15 of this study was compared to the ST15 outbreak lineage from Nepal (48, 49) by mapping against the completed genome of an isolate from the Nepal outbreak (49), which also carried four plasmids. The heatmap shows the mapping coverage across the chromosome before removal of recombination from white (low coverage) to black (high coverage), emphasizing the large deletions and acquisitions in the Pakistan and Nepal line, respectively. Even coverage of the genome could be observed, with three genetic islands missing in different subsets of our strains, indicating the dynamic within this closely related lineage. The tree was constructed after removing recombination by gubbins, bootstrap values are depicted as colour gradient; from red (0) to black (100). **(B)** A detailed view of two regions in the genome absent in groups of strains within the isolates from this study (plotted areas: 1473529-1493215, region 1; 3988478-4042608, region 2; coordinates from CP008929.1), but present in the lineage from Nepal, as well as other isolates from the same ST but not part of the clonal lineages as outgroups. In addition, the Nepal lineage acquired the chromosomal locus including yersiniabactin (23; plotted region 5155581-5264456); these three regions were also removed by gubbins before the tree construction.

The phylogeny indicated additional structure in the isolates from Lahore, which was confirmed by analysing the mapping results in more detail (Fig. 5). The mapping-based approach shows that, even within the tight cluster of the isolates sequenced in this analysis, at least three different patterns were identified. These altered patterns lack three regions found in the reference strain from Nepal (Fig. S7) and a detailed overview of resistance, virulence gene profiles, plasmid replicons and surface determinants (capsule and O-antigen type) is given in Fig. S6.

The first region is a gain in the Nepal lineage as it is also absent from more distantly related members of ST15, and covers the ICE element carrying yersiniabactin and other virulence determinants as described originally (Fig. S6, Fig. 5; 48, 49). The other two regions were selectively lost in parts of the ST15 lineage from the Lahore strain dataset (Fig. 5). Region 2, lost in 35 strains, includes a putative chitinase and cellulose synthase *acsAB*. In addition, this lineage has lost genes encoding type VI secretion systems, which are important for virulence as well as between-bacteria competition (50). We also note an additional type VI secretion locus located immediately downstream of the island is conserved. This region is recognised as an intact prophage by the phage prediction program phaster (51). Twenty of these strains have lost a third region that encoded genes involved in biofilm formation (*acsC*, *acsD*; the second copy of *bcsA* and *bcsB* required for cellulose biosynthesis; Fig. S6).

Only one of the Nepalese plasmids (plasmid A) is partially conserved in the isolates we examined (Fig. S7), in accordance with the absence of the genes for NDM-1 and yersiniabactin (Fig. S6). Several gene clusters from the Nepal outbreak plasmids are conserved and likely to be encoded on different plasmids, including (Nepal plasmid A) *lacIZY*, phosphate transport (*phnD*/*E ptxD*); arsenic (*ars*), copper (*pco*), silver (*sil*) and cation (*cus*) efflux systems, as well as plasmid maintenance proteins from this plasmid; only sparse similarity can be found with other plasmids (e.g. Nepal plasmid C streptomycin resistance), in accordance with the different virulence and resistance profiles of the two hospital lineages. A subgroup of the strains does not share the genes from plasmid A in Nepal, which can also be seen by their different virulence factor profile (Fig. S7).

## Discussion

The *Klebsiella* isolates represent routinely collected samples over a protracted period. We observed sporadic single-isolate lineages; smaller, related clusters of 5-10 isolates per lineage; in addition to two larger clusters of strains. One of the latter two represents members of *K. quasipneumoniae* subsp. *similipneumoniae*, a subspecies that was previously thought to be less virulent (Fig. 1; 15). The major *K. quasipneumoniae* sequence type (334) was so far observed in other studies as occasional isolate (e.g. 1/250 in 52; 1/167 in 53; 1/328 in 54; 2/198 in 55, whereas we report 52 (44 if excluding ST variants) isolates all originating from bloodstream infections, representing 29.2% (24.7%) of samples investigated. Given its similarity to *K. pneumoniae* in standard diagnostics, *K. quasipneumoniae* numbers could be potentially underestimated (56). *K. variicola*, which is equally difficult to distinguish from *K. pneumoniae*, occurs more frequently in clinical samples (e.g. 28/134 samples in 57; 9/198 in 55; 14/250 in 52; 20/328 in 54) and was suggested to be highly virulent (58), but we did not observe any strain belonging to this species.

With the continuous decline of effective antimicrobials and the lag in new ones being released, antibody-based treatments are becoming increasingly important and are being strongly supported by governments and global health programs (59, 60). There are challenges associated with immunoprophylaxis against *Klebsiella* infections as there is no single prevalent capsule- or O-antigen type; on the contrary, our data reveals a very high diversity in these potential vaccine targets especially with respect to O-Ag, that is generally far less variable than the capsule. A recent large-scale analysis of several *Klebsiella* datasets spanning a global collection identified serotypes O1, O2 and O3 in 80% of hospital infections (20); contrary to this previous finding, we observed a high number of non-O1/O2 types in our collection of *Klebsiella*; these are mainly, but not exclusively, contributed by the large number of *K. quasipneumoniae* (Fig. 1D, Table S1). This diversity, and the lack of apparent correlation to disease severity (20), indicate that switches to non-vaccine targeted serotypes might be easily done once immune pressure for certain types is applied and has been investigated in detail in other organisms (61, 62); a better understanding of the dynamics and diversity is crucial for the development of O-antigen based vaccines, which is ongoing.

Screening for the resistance and virulence genes revealed no single factors that characterise the most successful lineages or clinical outcome, which is different from other pathogens (e.g. 16, 18). Putative virulence factors in *Klebsiella* spp. are not well-defined beyond the polysaccharide capsule and typically include genes that are thought to mediate attachment, such as Mrk, or survival in the human host, e.g. siderophores (63). Despite spread and chromosomal integration of yersiniabactin however, aerolysin seem to be the main siderophore for *Klebsiella* with highest affinity for iron in *in vitro* experiments despite similar gene numbers expressed and its presence has a significant impact on virulence (64). The considerable variation in yersiniabactin systems suggests that there may be other selective pressures on iron acquisition systems such as antibodies or bacteriophage. The unique clinical manifestation of CG23 however might strongly depend on the unique feature of colibactin (i.e. enhanced gut colonisation and dissemination to the liver and brain; 65–67). Even though there are discussions whether hypervirulent and hypermucoid should be considered equally and the key virulence factor(s) is still not clear (25, 68, 69), the hallmarks are the virulence plasmid encoding the capsule regulator *rmpA/rmpA2*, aerobactin and metal resistances; and chromosomal integration of yersiniabactin and colibactin, microcin E942 and the sialic acid-containing serum-resistant capsule K1 (70). We report a strain set that is a hybrid type; either encoding the virulence plasmid but with *rmpA2* inactivated, or the yersiniabactin, but no strains with both features even though there are several variants circulating through the population. All strains with the virulence plasmid encode *rmpA2* as a pseudogene, and no strain in the collection encodes colibactin or the functional *rmpA*.

A similar observation was made on a set of recent strains, indicating that the accumulation of virulence factors is an ongoing process, and more work needs to be done of longitudinal studies to monitor the changes of the virulence potential of sequence types over time (71). Findings of these mixed strains, harbouring some but not all virulence factors known, emphasises the importance to understand if there is a negative fitness effect of the combination of all virulence factors or a mutual exclusivity of the mobile elements involved that are only made possible through the genetic background of CG23, or if virulence mechanisms simply spread somewhat slower, and we are only seeing a delay until they appear co-distributed with antimicrobial resistance plasmids.

The collection is derived from a time (2010 to 2012) where carbapenem resistance was still at comparatively low levels and only starting to spread, whilst it is at a very high prevalence today in *Klebsiella* in LMIC as well as high-income countries (1, 29, 31). The origin of NDM-1 is likely a gene fusion promoted by *Acinetobacter* (13), which was first reported in an IncA/C2 plasmid with the original cassette. Our dataset contains several lineages with NDM-1, however these did not become the major problems of the region in the subsequent years. We find three different plasmid constructs at least, with the classical IncA/C2 (similar to NZ_CP006661.1 = pNDM-US; HE016_3 found in *Klebsiella* 72), the IncHI2A/IncHI2 (NC_009838.1 = pAPEC-O1-R; H066_2 from *E. coli* 73), and a non-circularised (HE125) plasmid fragment, that showed highest similarity to a plasmid from *Klebsiella* and *E. coli* (KJ440075.1 = pLK78 74). Whilst the IncA/C2 and the MAR plasmid are nowadays spread widely through the *Klebsiella* population, the IncR plasmid is more diagnostic of MDR *E. coli*, and might have used *Klebsiella* rather as initial vehicle than a successful means of spread. The samples are from a time where ESBL strains were already widespread, and carbapenems used as optional treatment; an NDM-1 carrying plasmid would therefore provide a clear advantage at face value. However, the patterns are more complex and much remains to be understood about the interaction between plasmids and bacteria, but it seems clear that acquisition of an NDM-1 plasmid alone is neither a sign that the plasmid will now spread stably with the population, nor that this lineage has an undisputable advantage over all carbapenemase-negative lineages.

The genomics generally predicted the phenotype with respect to drug resistance. This is important as technology is being developed to identify resistance genes at the point-of-care. Subtle differences in the sensitivity that are overlooked by evaluating strains only below or above a specified drug concentration, and might indicate an additional mechanism to deactivate beta-lactamase inhibitors, especially since we could not detect any genomic differences between the beta-lactamases of closely related strains. Such an analysis does not preclude expression differences due to promoter differences that might impact phenotypic resistance. Monitoring this resistance in detail is important as piperacillin/tazobactam treatment is a common clinical response to ESBL infections (75). A recent report on stepwise resistance against an inhibitor also highlighted changes in the major porins (Omp35 and Omp36) as well as the LPS O-antigen locus (*rfb*); this could explain some resistances (e.g. the porin disruption in ST15) (76). AmpH is a penicillin-binding protein, but exhibits only extremely low beta-lactamase activity against nitrocefin (77, 78). None of the OXA alleles detected in our dataset conferred extended-spectrum beta-lactamase activity, but OXA enzymes are usually poorly inhibited by clavulanic acid, which might provide an important advantage (79).

Understanding the rapid spread of ESBL- and carbapenemase – carrying plasmids in combination with the spread of high-risk lineages arising from the diverse background population of *Klebsiella* is of crucial importance to the early identification of highly resistant and virulent clones, and preventing their spread. There is considerable subtlety in the evolution of drug resistance where having a variety of activity spectra across resistance enzymes is likely to be of importance, as enzymes with lower activity spectra are often more resistant to inhibitors than highly resistant ESBL enzymes (79). Detailed analyses are crucial to our understanding of the different mechanisms of resistance against alternative treatments such as beta-lactamase inhibitors. Inhibitors could become a highly effective treatment if resistance can be avoided, or more likely recognised earlier.

## Methods

### PacBio DNA preparation and sequencing

DNA was further purified by Phenol:Chloroform:Isoamyl Alcohol (25:24:1) and Chloroform:IAA (24:1) extractions using Phase-lock tubes (Qiagen) and re-dissolved in 10mM Tris pH7.4 buffer. Sequencing was performed on the PacBio RSII using P6/C4 sequencing chemistry, the library was prepared using the SMRTbell Template Prep Kit 1.0. Filtered sub-reads were generated with the pacbi-smrt software, and assembled with canu v1.1 (80). The assemblies were then circularised using circulator (81) with canu as assembler, and polished with unicycler-polish (82) and the Illumina reads of the respective sample. HE016 gave better results in assembling with the unicycler-hybrid assembler combining the PacBio data with the Illumina reads (15), and was subsequently circularised and polished as above. All assemblies were then annotated with prokka (83).

### Bacteria Mapping and Variant Detection

Mapping was performed against the chromosome of *Klebsiella pneumoniae* NTUH-K2044 (AP006725), a published whole genome from an outbreak in Nepal (49) and HE196 (this study). Sequence reads were mapped against the reference genome as indicated using SMALT (84; v0.7.4) to produce a BAM file. Variation detection was performed using samtools mpileup v0.1.19 bcftools v0.1.19 to produce a BCF file of all variant sites.

### Pan-genome analysis

The samples as indicated in the respective experiments were assembled and annotated as described above, and the GFF3 files generated by PROKKA v1.11 (83) were used as input for the pan-genome pipeline Roary (v3.7.0; 85) with a BLASTp percentage identity of 90% and a core definition of 99%. This gives a core gene alignment of 3486 genes for all strains from this study (Figure 1, 2). The core gene alignment was generated with mafft (86); SNPs were first extracted using snp-sites v2.3.2 (87), and then a maximum likelihood tree with RAxML (88) was calculated. For the ST15 comparison, the pan-genome was calculated as described above, and gene presence/absence analysed further using R and the resulting presence/absence matrix and phandango (89).

### Phylogenetic analyses

Trees were calculated using RAxML (v8.2.8; 88) with the time-reversible GTR model and 100 bootstrap repeats. Tree demonstrations were prepared in itol (90) as well as in house python and R scripts. For whole-genome mapping trees, recombinant regions were removed using Gubbins (v1.4.9; 91) and a maximum likelihood tree calculated with RAxML to obtain bootstrap support values, as described above, and sites with more than 5% N were not considered in the tree calculation.

### Gene content analysis

For the determination of the antibiotic resistance profile, the virulence factor profile and plasmid types, we used ariba The resistance, virulence and plasmid profiles were matched against the modified version of ARG-ANNOT (92) available at the SRST2 website (https://github.com/katholt/srst2/tree/master/data; download date 02.10.2016), a dataset of virulence factors obtained from the *Klebsiella*-specific BIGSDB (http://bigsdb.pasteur.fr/klebsiella/klebsiella.html; download date 22. 02. 2016), and the PlasmidFinder database as implemented in ariba v2.10.0 (93, 94). Sequence gene profiles and types were determined using MLST check (95) comparing assembled genomes against the MLST database for *Klebsiella pneumoniae* (pubmlst.org/Klebsiellapneumoniae/). For further resistance determinants (SNP-based or porin inactivation), a database with the genes of interest was created as outlined in the ariba manual pages. The assignment of *bla-SHV* alleles was controlled based on amino acids using the beta-lactamase database http://www.laced.uni-stuttgart.de. Yersiniabactin and ICE alleles were annotated using kleborate (23; https://github.com/katholt/Kleborate). Capsule and LPS O-antigen was annotated using kleborate and a custom database of LPS O-antigens (20). The assignment to O3 subgroups was updated based on recent nomenclature (O3l/s; now a/b/c) using the online tool captive (http://kaptive.holtlab.net/, 22).

### Minimum inhibitory concentrations (MIC) measurements

All isolates were analysed using the VITEK 2 system (bioMérieux, UK). In brief, suspensions of colonies were made in 0.45% saline solution from growth on iso-sensitest agar (Thermo Scientific Oxiod, UK), adjusted to a turbidity equivalent to that of a 0.5 McFarland standard and used to load the test cards, using manufacturer’s instructions. The *Enterobacteriaceae* AST-N350 cards was automatically filled, sealed and inserted into the VITEK 2 reader–incubator module (incubation temperature 37°C), and fluorescence measurements were performed every 15 min for up to 18 h. For detailed analysis of several NDM-1 containing strains, the MICs for meropenem were also assessed manually following the protocol of Wiegand 2008 (96; Table 1).

## Data availability

Details on strains and accession numbers can be found in Table S1. All relevant data is in the manuscript and supporting material, alignment and tree files are available for download at figshare https://figshare.com/s/cdaddb659a6e178102df.

**Supplement: Figures S1-S7 (with Figure legends) and Tables S1-S3:**

**Table S1: Details on the strains from Pakistan.**

**Table S2: Vitek measurements.**

**Table S3: Additional strains included in the detailed analysis of ST15.**

## Acknowledgments

The authors thank Karen Oliver for help with the PacBio sequencing, and Andrew Page for helpful discussions on PacBio assembly strategies, as well as the Pathogens Informatics group. We thank Dr. Badr Alzahrani (Jouf University) for comments on the manuscript. This work was supported by the NHMRC (Program Grant 1092262), the Wellcome Trust (206194), and the Higher Education Commission of Pakistan and The Children’s Hospital & The ICH, Lahore, Pakistan. H.E. was supported by a scholarship from Higher Education Commission (HEC) Pakistan under the International Research Support Initiative Program (IRSIP).

