## Supplementary Materials for "Emergence of carbapenem, beta-lactamase inhibitor and cefoxitin resistant lineages from a background of ESBL-producing *Klebsiella pneumoniae* and *K. quasipneumoniae* highlights different evolutionary mechanisms"

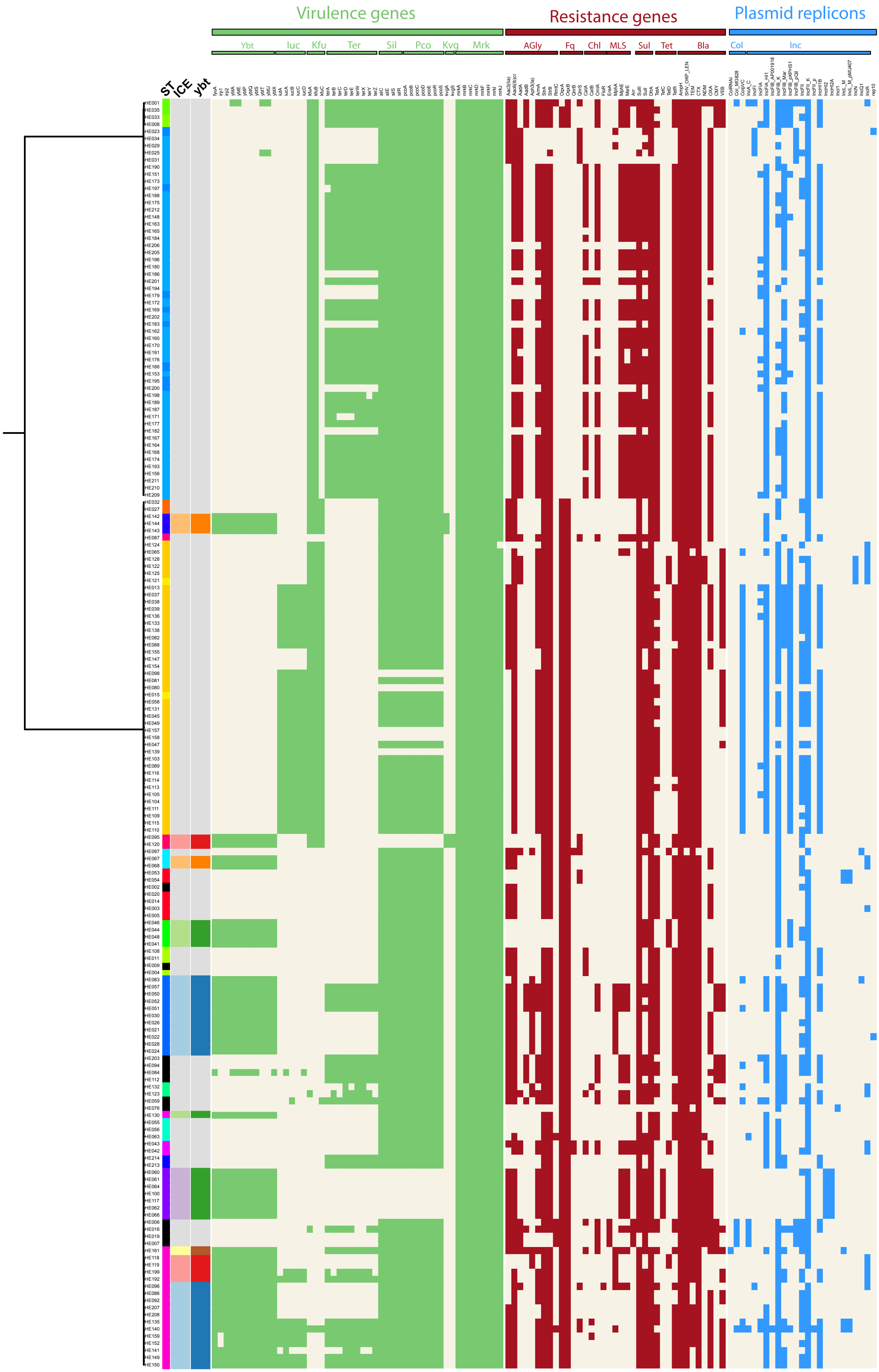

Virulence genes
Resistance genes
Plasmid replicons
Ybt
Iuc
Kfu
Ter
Sil
Pco
Kvg
Mrk
AGly
Fq
Chl
MLS
Sul
Tet
Bla
Col
Inc
ST
ICE
ybt

**Fig. S1: Antimicrobial resistance (AMR), plasmid replicons and virulence genes.** The predictions of AMR genes, virulence genes and plasmid replicons was performed using ariba (93), a mapping-based approach independent of assemblies. The guidance tree is based on the core gene alignment for this strain set generated by roary (85).

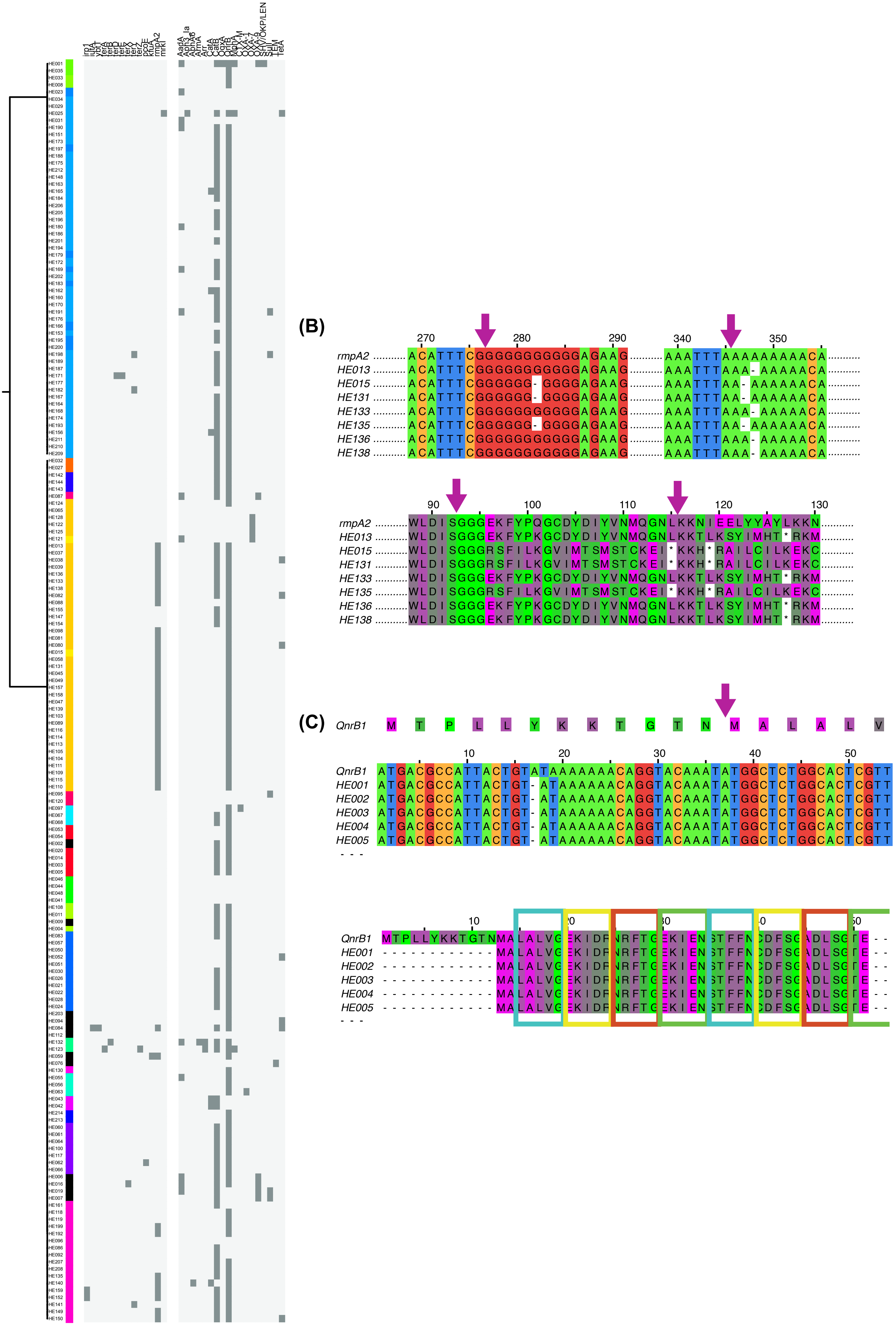
(B)
270 280 290 340 350
rmpA2 ACATTTCGGGGGGGGGGGAGAAG AAATTTAAAAAAAAACA
HE013 ACATTTCGGGGGGGGGGGAGAAG AAATTTAAA-AAAAACA
HE015 ACATTTCGGGGGG-GGGGAGAAG AAATTTAA-AAAAAACA
HE131 ACATTTCGGGGGG-GGGGAGAAG AAATTTAA-AAAAAACA
HE133 ACATTTCGGGGGGGGGGGAGAAG AAATTTAAA-AAAAACA
HE135 ACATTTCGGGGGG-GGGGAGAAG AAATTTAA-AAAAAACA
HE136 ACATTTCGGGGGGGGGGGAGAAG AAATTTAAA-AAAAACA
HE138 ACATTTCGGGGGGGGGGGAGAAG AAATTTAAA-AAAAACA
90 100 110 120 130
rmpA2 WLDISGGGEKFYPQGCDYDIYVNMQGNLKKNIEELYYAYLKKN
HE013 WLDISGGGEKFYPKGCDYDIYVNMQGNLKKTLKSYIMHT*RKM
HE015 WLDISGGGRSFILKGVIMTSMSTCKEI*KKH*RAILCILKEKC
HE131 WLDISGGGRSFILKGVIMTSMSTCKEI*KKH*RAILCILKEKC
HE133 WLDISGGGEKFYPKGCDYDIYVNMQGNLKKTLKSYIMHT*RKM
HE135 WLDISGGGRSFILKGVIMTSMSTCKEI*KKH*RAILCILKEKC
HE136 WLDISGGGEKFYPKGCDYDIYVNMQGNLKKTLKSYIMHT*RKM
HE138 WLDISGGGEKFYPKGCDYDIYVNMQGNLKKTLKSYIMHT*RKM
(C)
QnrB1 M T P L L Y K K T G T N M A L A L V
10 20 30 40 50
QnrB1 ATGACGCCATTACTGTATAAAAAAACAGGTACAAATATGGCTCTGGCACTCGTT
HE001 ATGACGCCATTACTGT-ATAAAAAAACAGGTACAAATATGGCTCTGGCACTCGTT
HE002 ATGACGCCATTACTGT-ATAAAAAAACAGGTACAAATATGGCTCTGGCACTCGTT
HE003 ATGACGCCATTACTGT-ATAAAAAAACAGGTACAAATATGGCTCTGGCACTCGTT
HE004 ATGACGCCATTACTGT-ATAAAAAAACAGGTACAAATATGGCTCTGGCACTCGTT
HE005 ATGACGCCATTACTGT-ATAAAAAAACAGGTACAAATATGGCTCTGGCACTCGTT
- - -
QnrB1 MTPLLYKKTGTNMALALVGEKIDRNRFTGEKIENSTFFNCDFSGADLSGTE- -
HE001 - - - - - - - - - - - - MALALVGEKIDRNRFTGEKIENSTFFNCDFSGADLSGTE- -
HE002 - - - - - - - - - - - - MALALVGEKIDRNRFTGEKIENSTFFNCDFSGADLSGTE- -
HE003 - - - - - - - - - - - - MALALVGEKIDRNRFTGEKIENSTFFNCDFSGADLSGTE- -
HE004 - - - - - - - - - - - - MALALVGEKIDRNRFTGEKIENSTFFNCDFSGADLSGTE- -
HE005 - - - - - - - - - - - - MALALVGEKIDRNRFTGEKIENSTFFNCDFSGADLSGTE- -
- - -

**Fig. S2: Predicted virulence and resistance genes with lost function. (A)** Core gene tree as in Fig. 1, the heatmap indicates genes that were predicted as pseudogenes by ariba due to missing start and/or stop codons or presence of a stop codon interrupting the sequence. **(B)** The sequences retrieved from ariba predicted as *rmpA2* pseudogenes were retrieved and aligned to a reference sequence (amino acids NP_943354.1 and the respective CDS). Two homopolymer regions, a poly-G and a poly-A repeat, show mutations that lead to subsequent frameshifts and stop codons in the translated sequence. The arrows mark the corresponding positions in the amino acid and nucleotide alignments to facilitate orientation. Colours, nucleotides (upper panel); helix propensity (lower panel); as implemented in JalView (99). **(C)** The N-terminal region of *qnrB* encodes a frameshift in our data, however an alternative start codon can be recognised (arrow), and leads to a full-length product including all structural elements as in the structure (34). Colours upper panel, nucleotides; colours lower panel, helix propensity; as implemented in JalView (99). Structural elements shown as coloured boxes as in Figure A1 from (34).

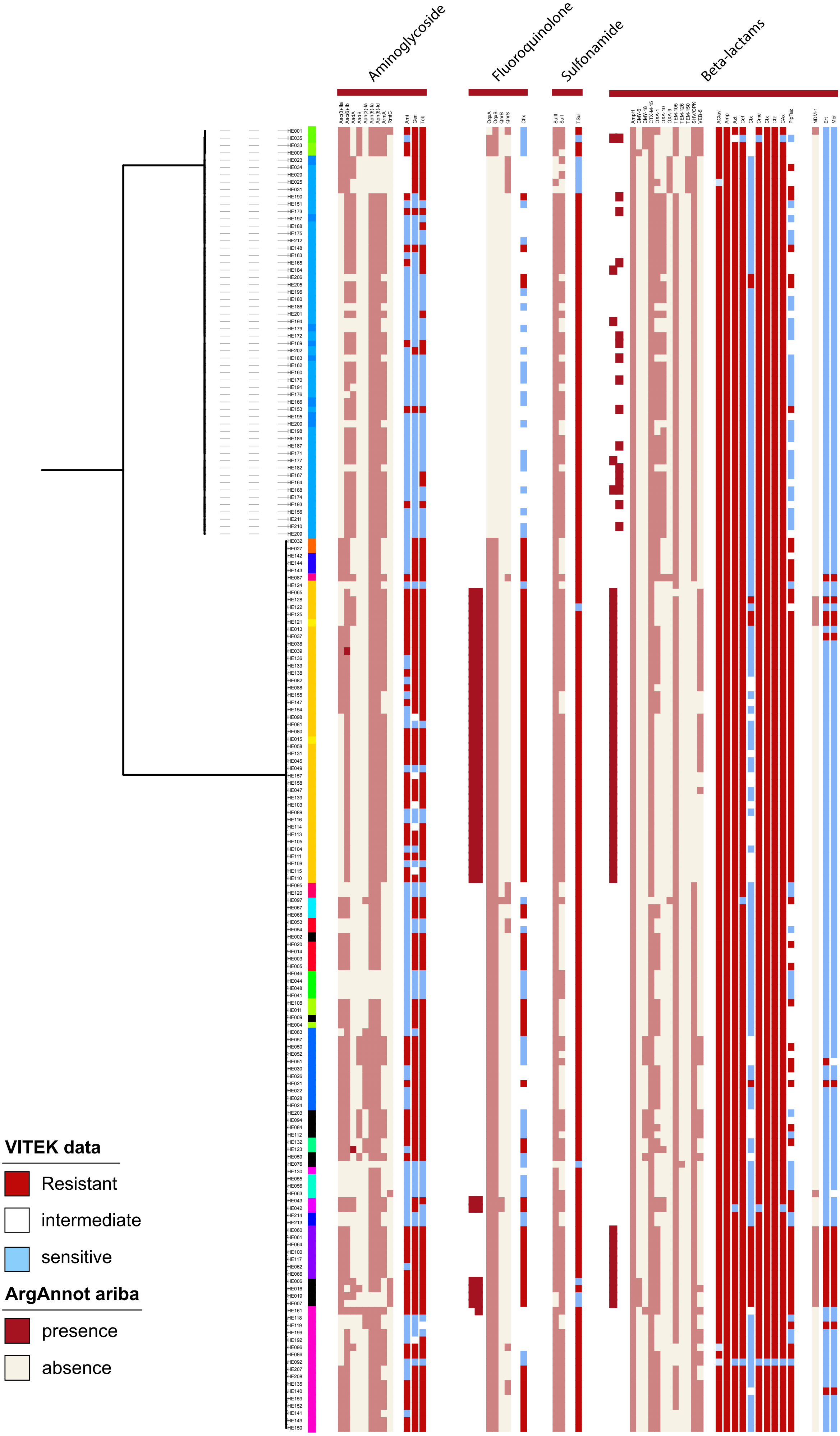
Aminoglycoside
Fluoroquinolone
Sulfonamide
Beta-lactams
VITEK data
Resistant
intermediate
sensitive
ArgAnnot ariba
presence
absence

**Fig. S3: Comparing measured minimum inhibitory concentration (MIC) data with predicted resistance types.** Vitek measurements are compared with the predicted known resistance-conferring genes for aminoglycosides, fluoroquinolones, sulfonamindes and beta-lactams (including carbapenems). Red indicates resistant, white indicates intermediate, blue indicates sensitive measurement.

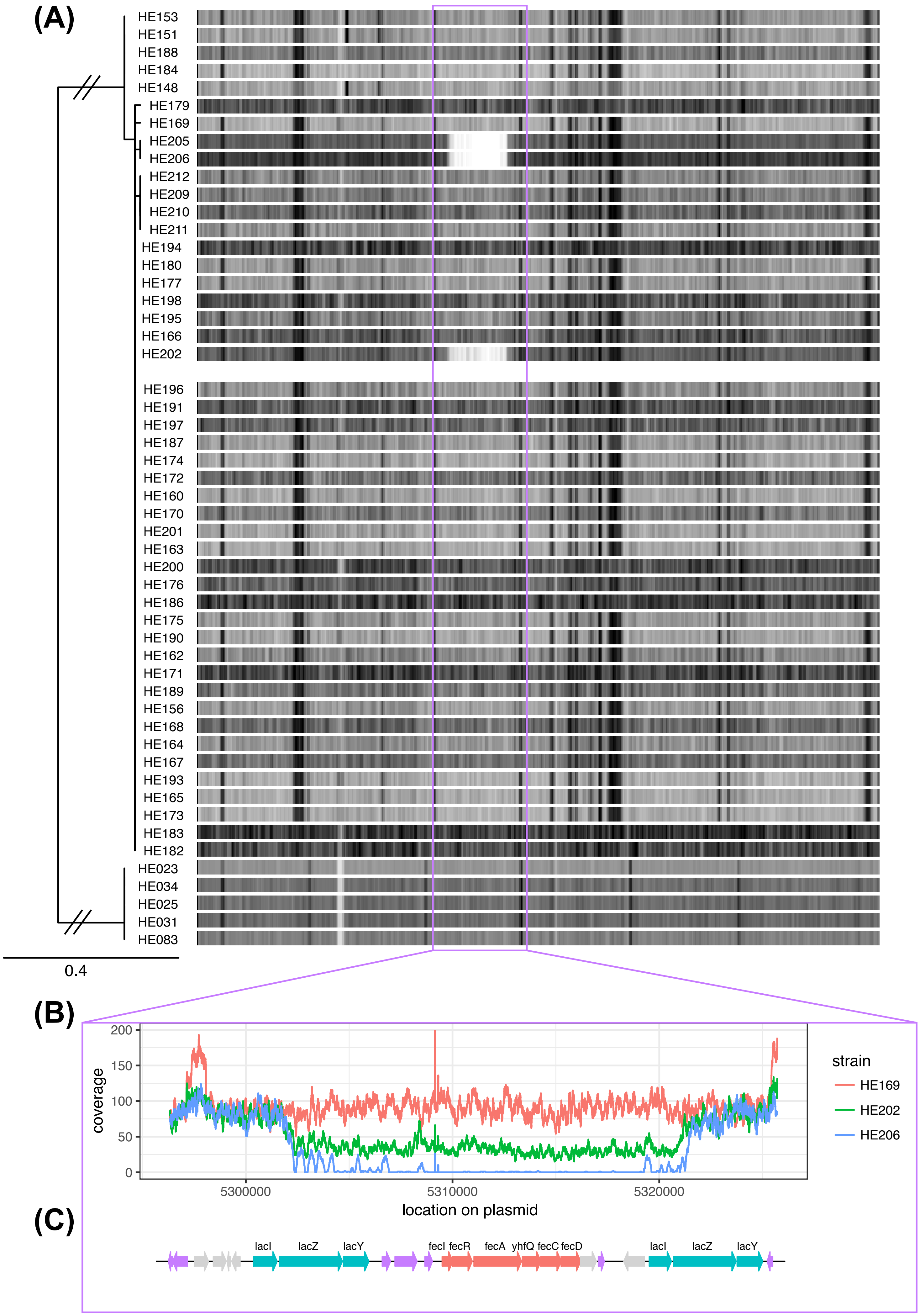
(A)
HE153
HE151
HE188
HE184
HE148
HE179
HE169
HE205
HE206
HE212
HE209
HE210
HE211
HE194
HE180
HE177
HE198
HE195
HE166
HE202
HE196
HE191
HE197
HE187
HE174
HE172
HE160
HE170
HE201
HE163
HE200
HE176
HE186
HE175
HE190
HE162
HE171
HE189
HE156
HE168
HE164
HE167
HE193
HE165
HE173
HE183
HE182
HE023
HE034
HE025
HE031
HE083
0.4
(B)
200
150
100
50
0
coverage
5300000
5310000
5320000
location on plasmid
strain
HE169
HE202
HE206
(C)
lacI
lacZ
lacY
fecI
fecR
fecA
yhfQ
fecC
fecD
lacI
lacZ
lacY

**Fig. S4: Conservation of the plasmid in *K. quasipneumoniae* ST334 isolates. (A)** Mapping tree against the chromosomal contig of HE196 resolved by PacBio; the heatmap shows the mapping density against the plasmid HE196-2. **(B)** Detailed coverage of the region missing in two, and only low-coverage in one of the strains (5296336-5325706 on the reference sequence), as well as a schematic of the respective predicted genes for the region.

(A)

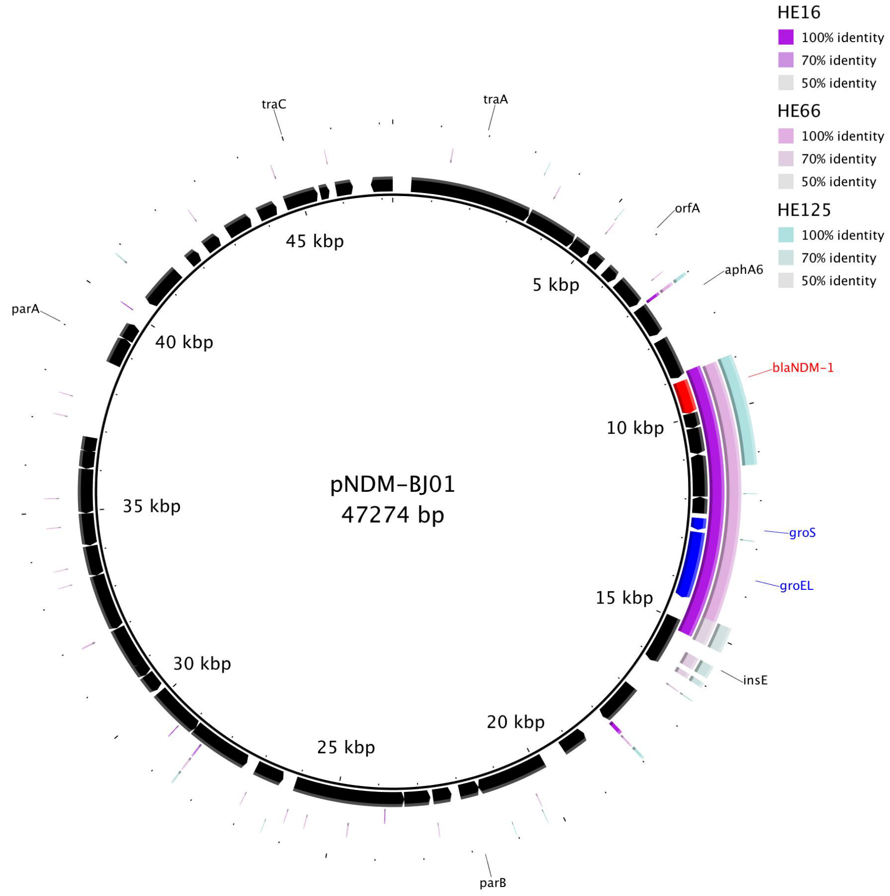
HE16
100% identity
70% identity
50% identity
HE66
100% identity
70% identity
50% identity
HE125
100% identity
70% identity
50% identity
traC
traA
orfA
aphA6
blaNDM-1
groS
groEL
insE
parB
parA
5 kbp
10 kbp
15 kbp
20 kbp
25 kbp
30 kbp
35 kbp
40 kbp
45 kbp
pNDM-BJ01
47274 bp

(B)

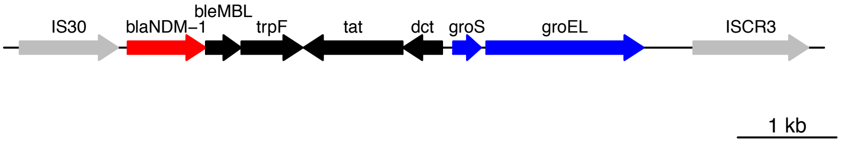
IS30
blaNDM-1
bleMBL
trpF
tat
dct
groS
groEL
ISCR3
1 kb

**Fig. S5: Conservation of the original NDM-1 cassette.** (A) The three plasmid contigs from the PacBio assemblies of HE016, HE066 and HE125 which contain the NDM-1 gene, were compared against a plasmid encoding the original cassette as described for *Acinetobacter* (100). This shows the different levels of conservation of the cassette, whilst nothing else of the original plasmid can be found; **(B)** shows a close-up of the original NDM-1 mobile element (13).

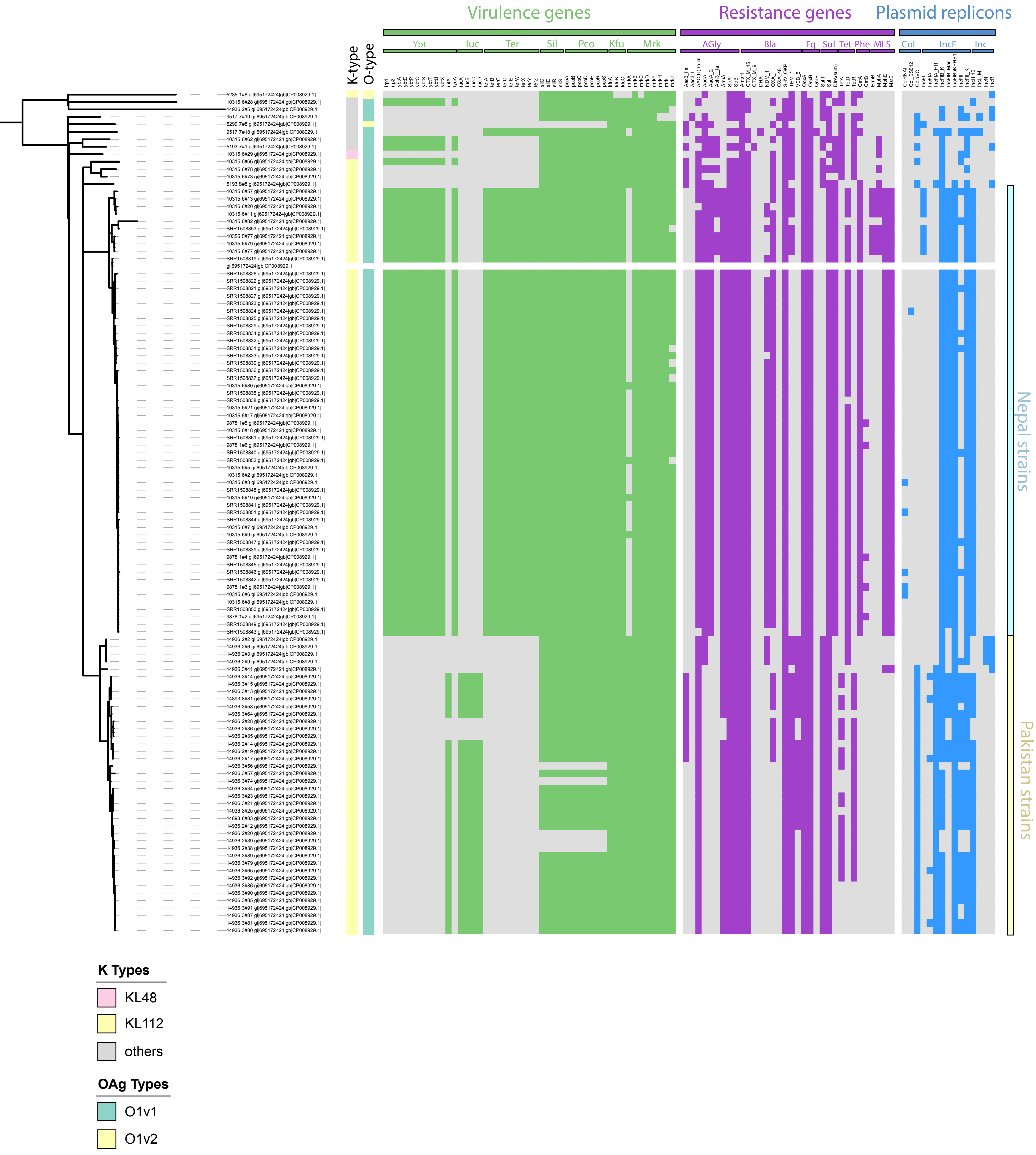
Virulence genes
Resistance genes
Plasmid replicons
Ybt
Iuc
Ter
Sil
Pco
Kfu
Mrk
AGly
Bla
Fq
Sul
Tet
Phe
MLS
Col
IncF
Inc
K-type
O-type
Nepal strains
Pakistan strains
K Types
KL48
KL112
others
OAg Types
O1v1
O1v2

**Fig. S6: Comparison of the ST15 isolates from an outbreak in Nepal and our collection from Pakistan.** Plasmid replicons, resistance genes and virulence genes were predicted by ariba; the tree is a whole-genome tree after removing recombination with gubbins against the reference strain from Nepal (CP008929.1).

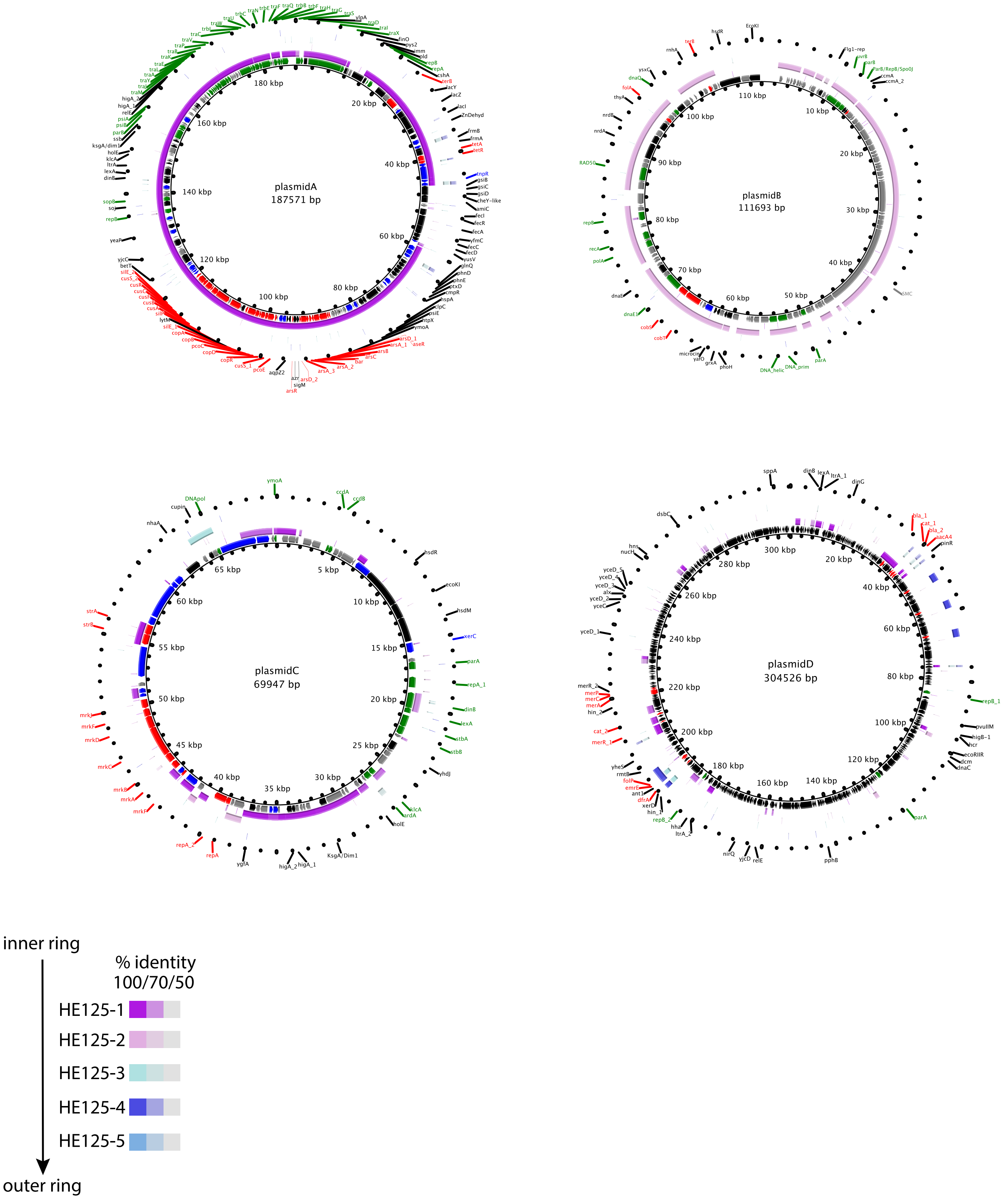
plasmidA
187571 bp
plasmidB
111693 bp
plasmidC
69947 bp
plasmidD
304526 bp
inner ring
outer ring
% identity
100/70/50
HE125-1
HE125-2
HE125-3
HE125-4
HE125-5

**Fig. S7: Comparison of the plasmid profiles between the Pakistan and Nepal ST15 lineage.** The full genome sequence of a representative strain from the Nepal outbreak (49) was used as reference, and all five plasmid contigs from HE125, a representative of the related ST15 lineage from Nepal, were compared against the four plasmids as annotated (plasmid A = CP008930.1, B = CP008931.1, C = CP008932.1, D = CP008933.1). The legend indicates similarity values for the different plasmid contigs from the HE125 strain.
